# Air leaks: stapling affects porcine lungs biomechanics

**DOI:** 10.1101/2021.04.19.433417

**Authors:** Bénédicte Bonnet, Ilyass Tabiai, George Rakovich, Frédérick P. Gosselin, Isabelle Villemure

**Author notes:** **Corresponding author**: Isabelle Villemure.

## Abstract

During thoracic operations, surgical staplers resect cancerous tumors and seal the spared lung. However, post-operative air leaks are undesirable clinical consequences: staple legs wound lung tissue. Subsequent to this trauma, air leaks from lung tissue into the pleural space. This affects the lung’s physiology and patients’ recovery.

The objective is to biomechanically and visually characterize porcine lung tissue with and without staples in order to gain knowledge on air leakage following pulmonary resection. Therefore, a syringe pump filled with air inflates and deflates eleven porcine lungs cyclically without exceeding 10 cmH_2_O of pressure. Cameras capture stereo-images of the deformed lung surface at regular intervals while a microcontroller simultaneously records the alveolar pressure and the volume of air pumped. The raw images are then used to compute tri-dimensional displacements and strains with the Digital Image Correlation method (DIC).

Air bubbles originated at staple holes of inner row from exposed porcine lung tissue due to torn pleural on costal surface. Compared during inflation, left upper or lower lobe resections have similar compliance (slope of the pressure vs volume curve), which are 9% lower than healthy lung compliance. However, lower lobes statistically burst at lower pressures than upper lobes (*p-*value<0.046) in *ex vivo* conditions confirming previous clinical *in vivo* studies. In parallel, the lung deformed mostly in the vicinity of staple holes and presented maximum shear strain near the observed leak location. To conclude, a novel technique DIC provided concrete evidence of the post-operative air leaks biomechanics. Further studies could investigate causal relationships between the mechanical parameters and the development of an air leak.

## 1. Introduction

To treat patients with lung cancer, thoracic surgeons remove the cancerous tumor. This resection surgery cuts out a segment, a lobe or several lobes to a whole lung. To do so, a surgical stapler squeezes tissue around the tumor to cut and seal the healthy tissue from the tumor bearing one. After chest surgery, 28-60% of patients suffer from air leaks [1]– [4]. Patients routinely require a chest tube (drain) until the leak resorbs [2], [5], which results in longer hospital stays and increased care costs [6].

A postoperative air leak was defined by Mueller et. al. [2] as “air escaping the lung parenchyma into the pleural space after any kind of surgery in the chest”. However, very few *ex vivo* studies described the origin of the leak [2], [4], [7]–[10]. Imhoff *et. al*. [7] resected 18 dog lung lobes with a surgical stapler, 3 cm away from their edges while being under 10 cmH2O of pressure and further immersed them to observe the leak. They stated that leak is observed with bubbles at the staple holes and at the extremities of the staple line [8]. The main limitations of this study are that: different lobes were tested regardless of whether they came from the left or right lung nor their size; the lobes were inflated during stapling as opposed to normal surgical stapling conditions where the lung is deflated, which may contribute to poor sealing at the staple line; the pressure was increased instead of being cycled to better mimic ventilation conditions; they mentioned that the leak happened at staple lines but did not provide evidence for this claim [9], [11], [12].

At the clinical level, data suggest that the incidence of air leaks varies with the type, extent and location of the resection [13]–[15]. In particular, air leak risks increase with upper lobectomies [16]–[18]. This last statement was investigated in the only available numerical study done by Casha *et. al*. [1]. They concluded that lower lobes leak most because they need to reshape to fit the rib cage bullet-shape exposing them to higher mechanical stress. Other clinical studies explored the consequences of stapling. Resections were found to reduce the respiratory system compliance *C* = ΔV/ΔP (mL/cmH_2_O), decreasing the lung’s ability to deform with the rib cage [19]–[21]. Finally, short time post-operative air leaks supposedly originate from biomechanical causes, suggesting that they are related to staple and mechanical parameters such as pressure and volume [22]. Considering the field of air leaks after lung surgery, there are very few published fundamental research studies investigating the biomechanics of the problem [7], [8], [23], [24]. Recently, Digital Image Correlation (DIC) has been a technique of choice for accurately measuring the strain on the surface of biological materials [24]– [26]. To our knowledge, DIC is a novel approach for studying interactions between pleura-covered lung tissue and staples at staple or whole organ scale.

Based on these published findings, we have hypothesized as follows: the air leakage would be the result of the development of a crack in the pleura, due to its perforation by the staple legs, exposing lung tissue that would then not prevent air from escaping. Taken together, these studies highlight the importance of investigating the biomechanics of the stapling-induced leakage. Therefore, this article tackles the following question: how do the mechanical parameters of the “lung tissue/staples” complex, i.e. volume, pressure and strain, influence air leakage in porcine lung resections with surgical staplers? The purpose of this paper is to biomechanically and visually characterize porcine lung tissue with and without staples in order to gain knowledge on air leakage following pulmonary resection. The next sections of this paper present a description of the experimental methods, including the positive pressure ventilation mechanism, the optical method for strain fields measurements, results analyses and statistical tests, the qualitative and quantitative results concerning air leakage, a discussion section including limitations, and a conclusion.

## 2. Material and methods

This section describes the experimental setup used to ventilate the lung and monitor its volume and pressure as well as the optical method that measures the strain of the lung surface. The protocol described below was approved by Research Department of Polytechnique Montreal (BIO-1819-09) prior to experiments.

### 2.1. Tissue preparation

Porcine red offal was obtained immediately after the death of the animals from a local slaughterhouse. Pig lungs were chosen for their anatomic resemblance to human lungs and ease of procurement. Among this offal, thirty-two lungs from medium to large pigs stored in a cooler at 4°C were received. To limit the study parameters, the left lung, which has only two lobes, was selected to simulate two surgeries (section 2.4). Unlike the right lung, the bronchial structure of the left lung allows straightforward access for ventilation. The condition of the left lung was visually checked and any specimen showing a scratch of the visceral pleura or lung tissue was rejected. The tissue was then prepared for the experiment, carried out within 48 hours of the animal’s sacrifice.

Specimens were stored at 4°C to preserve their mechanical properties [27], [28]. A scalpel separated the left lung from the rest of the lung complex (trachea, right lung, pulmonary ligament) and a scale weighed the sample. The lobar bronchi were dissected from the lung parenchyma. A cyanoacrylate glue bonded the flexible PVC tube connected to the syringe pump to the left main bronchus (Figure 1). The left lung was positioned on a stainless-steel table protected by two layers of protective absorbent paper and a layer of aluminum foil. This allowed the lung to freely move while protecting the material from biological contamination. The mediastinal side rested on the table.

**Figure 1:**
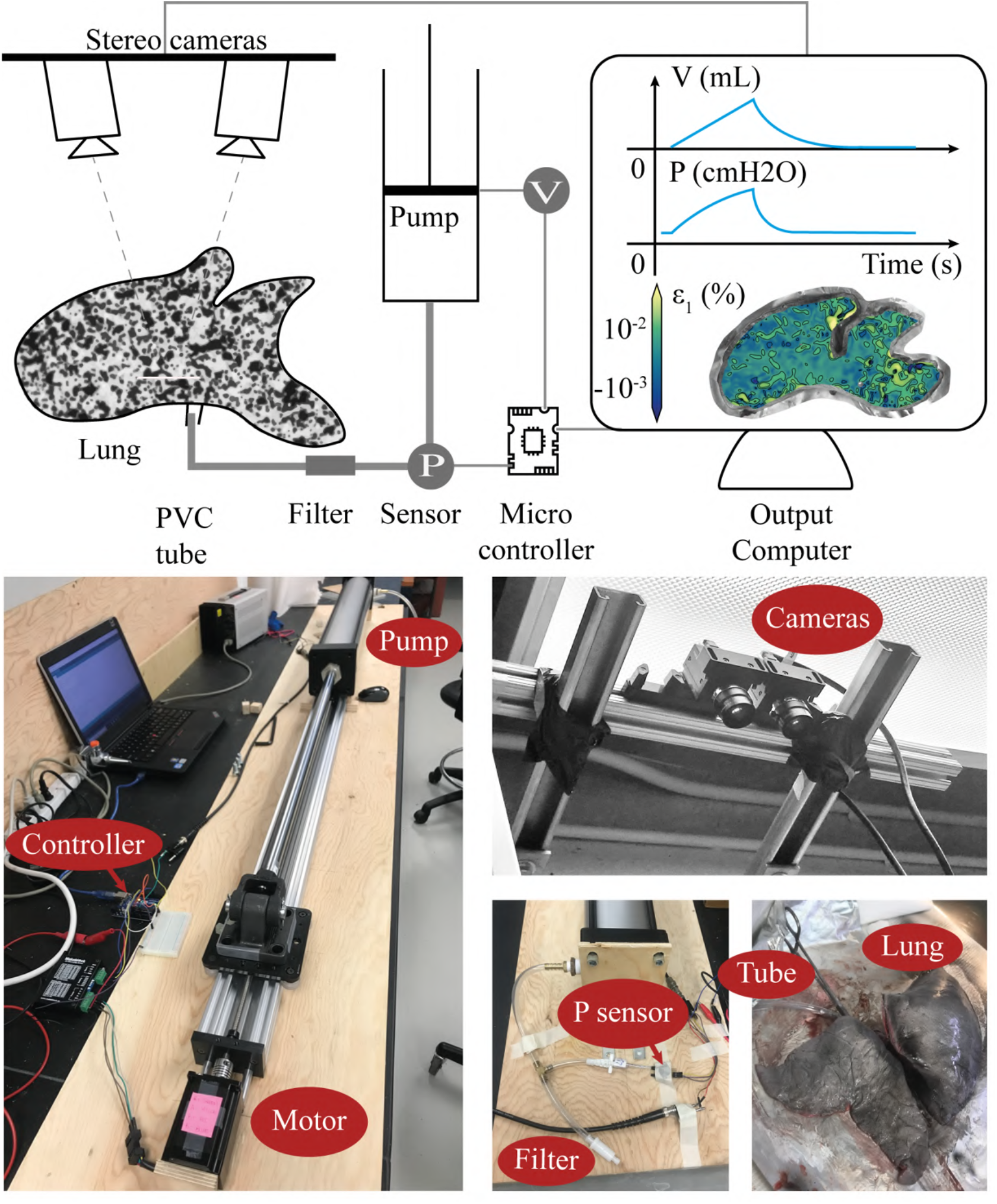
Experimental setup. Arduino-controlled syringe-pump delivers air to a left porcine lung passing through an in-line filter via a PVC flexible tubing. Arduino microcontroller collects pressure measurements from a pressure sensor and volume measurements from piston position. Two cameras record from above the 3D curved costal surface of the lung. The screen shows an example the pressure and volume versus time curves recorded by the microcontroller as well as the strain patterns obtained by DIC. SIZE: 1.5 COLUMNS-FITTING

### 2.2. Experimental setup

A microcontroller (Arduino UNO, Figure 1) controlled a linear actuator (C-Beam® XLarge Linear Actuator Bundle, OpenBuilds) assembled with a cylinder (30” long, 4” diameter). A flexible PVC tubing (Figure 1) connected the pump (Figure 1) to the lung passing through an in-line filter (Figure 1) that prevented the contamination of the material with biological particles. A calibrated differential pressure sensor (MPX5010DP, NXP, Figure 1) measured the air pressure delivered by the syringe pump relative to the atmospheric pressure, which was equal to the alveolar pressure in quasi-static conditions (5 mL/s). This measurement was made downstream of the syringe pump before the air reached the lung (Figure 1) through the flexible PVC tube. A coaxial cable transmitted the measured pressure values to the microcontroller and the acquisition board in analog signal. The Arduino controlled air supply (Figure 1) from or to the lung as a function of the alveolar pressure of the lung (Figure 1). The Arduino acquires volume and pressure data at 30 Hz. The custom designed syringe pump therefore inflated and deflated the porcine lung to simulate positive-pressure ventilation.

Two 5 Megapixels Pointgrey™ cameras (acquired from Correlated Solutions Inc.) were attached over the experimental setup in order to observe the curved costal surface of the left lung as seen from above. The field of view was aligned with the center of the specimen. To prepare the lung surface for DIC, an absorbent paper pad first dried out the curved costal surface of the left lung. A first layer of white foundation (SnazarooTM, UK) covered this studied portion of the specimen at rested state. A second layer was applied while the lung was inflated at a pressure of 10 cmH_2_O. Then, an airbrush (0.2 mm nozzle diameter, 0.2 mm needle diameter, GocheerTM, China) painted on this white background a random pattern of black dots with a black water-based paint (Golden Artist Colors, Inc., USA) called a speckle. Several diffuse light sources were chosen to illuminate the specimen homogeneously while minimizing reflections.

The VIC-Snap software (Correlated Solutions Inc., South Carolina, Figure 1) gave real-time feedback on the quality of the contrast, focus and lighting of the setup. The DIC method compared a subdivision (section 2.5) taken from the reference image (at resting state) to all the subdivisions of the deformed image until the subdivision of the deformed image matched the reference subdivision. It also estimated the quality of the speckle pattern and experimental setup with the correlation error (one standard-deviation confidence interval), before acquiring any data. Contrast, focus and lighting were adjusted to obtain the lowest correlation error. A calibration plate of 20.9 mm x 15.3 mm with dot spacing of 14 mm was used to calibrate the DIC-system. The VIC-Snap software controlled image acquisition at 3 Hz by the cameras. At the same time, the software saved pressure values sent to the acquisition board by the pressure sensor. The DIC captured the displacements of the black speckle pattern during the test (Figure 1) and thereafter analysis computed the strain patterns of the costal surface of the lung.

### 2.3. Ventilation procedure

Once the left lung was prepared with its speckle pattern, it was placed under the DIC cameras. To restore the mechanical equilibrium of the lung (P_sensor_ = 0 cmH_2_O), the specimen was disconnected from the syringe pump at the end of each experiment and reconnected before a new experiment begins. Lung preconditioning (Figure 2) consisted of 10 breathing cycles, including one inhalation followed by one exhalation per cycle, between pressures of 2 and 5 cmH_2_O. Positive pressure ventilation was used to reproduce surgical ventilation conditions of the lungs. Pressure controlled ventilation was preferred because the animal’s weight was not precisely known and necessary to assess the volume of a breath to perform volume controlled ventilation. The positive end-expiratory pressure was chosen to avoid atelectasis and maintain alveoli open as they may collapse at lower pressure than 2 cmH_2_O. The peak inspiratory pressure (PIP) was first set at 5 cmH_2_O for removal of residual atelectasis. The respiration flow was constant at 5 mL/s during inhalation and at 2.5 mL/s during exhalation, followed by a 5 second pause before the next cycle, approximating physiological conditions [29]. The healthy lung subsequently underwent three different tests, in which the PIP was incremented from 5 cmH_2_O to 10 cmH_2_O as follows: 10 breathing cycles between 2 and 5 cmH_2_O, 10 breathing cycles between 2 and 7 cmH_2_O, 10 breathing cycles between 2 and 10 cmH_2_O (Figure 2). The chosen PIP values were set to remain within the linear portion of the compliance curve estimated from preliminary results. It also allowed to measure lung compliance and to ensure that the displacement magnitude remained within the DIC working span (< 10 %) to avoid the loss of correlation in DIC measurements. In addition, it preserved the integrity of staple lines because an air leak at this point in the experiment would have made it impossible to obtain intended measurements.

**Figure 2:**
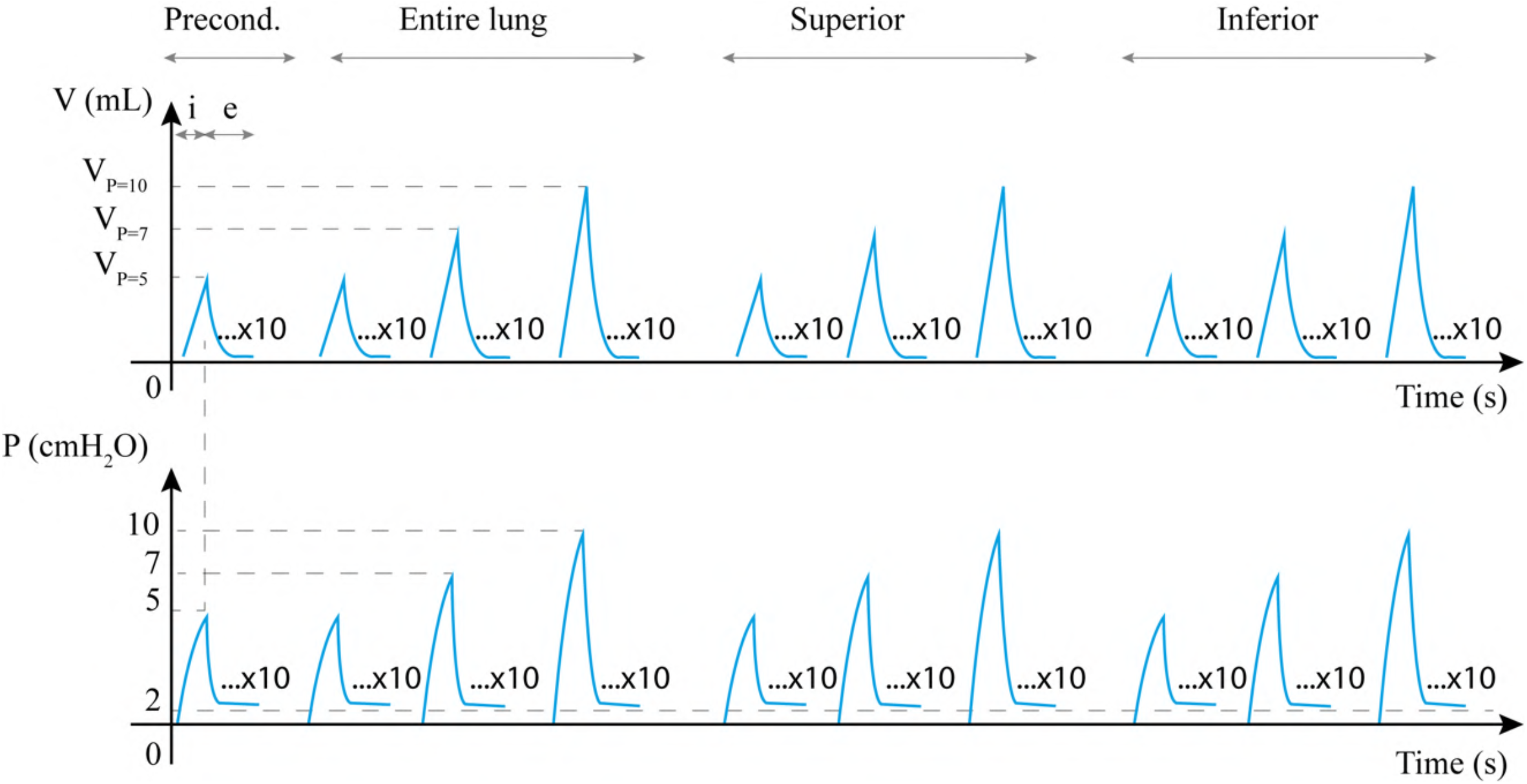
Schematics of the testing procedure. Volume (mL) and pressure (cmH_2_O) curves as a function of time for preconditioning, ventilation of the entire healthy lung, ventilation of the superior lobe and of inferior lobe alone. Each ventilation, including inflation followed by exhalation, is cycled 10 times. SIZE: 1.5 COLUMNS-FITTING

Then, the healthy lung underwent the resection surgery (section 2.4). The same testing procedure (Figure 2) was applied separately to the lower and upper lobes. If a portion of the speckle pattern was altered by the resection surgery, then it was redone completely to allow proper measurements with DIC. Finally, each lobe was inflated separately up to 60 cmH_2_O to replicate high physiological pressure and measure the burst pressure of the staple line identified by a change in volume or reaching a pressure plateau during ventilation. Once the test procedure was completed, the lung tissue was cleaned with water. A compressed air source then inflated the lung in a water tank to visually observe air bubbles, thus identifying the specificity of location of the air leak.

### 2.4. Resection procedure

A thoracic surgeon trained the experimenter to perform a division of the fissure between the lobes while isolating the uninjured lobar bronchi. This procedure [30] uses an EndoGIA™ linear surgical stapler and two green universal cartridge reloads (length 60 mm – width 4.8 mm, EndoGIA™, Medtronic) obtained from a product donation of Medtronic (Medical and Regulatory Affairs, Canada). This choice being very much surgeon dependent, the cartridge size was fixed so it would not become a parameter of the study. Each cartridge consists of two sets of three staggered rows of staples positioned on either side of a linear guide that is traversed by the stapler’s cutter. During the surgery, a small caliber rubber tube guided the anvil of the stapler atraumatically between the bronchial bifurcation and the lung tissue. The stapler was then fired to divide the lung parenchyma along the interlobar fissure. Staples were fired on the costal surface of the lung, from the outside toward the bronchus completing the natural separation (fissure) of the lung to independently separate the left upper lobe from the left lower lobe. After stapling, each lobe is sealed with a set of three rows of staples called a staple line. The inner row is in contact with the lung tissue, which swells when breathing, while the furthest row is close to the cutting line made by the cutter. Although two cartridges usually sufficed to divide the entire fissure, the last few millimeters of tissue were stapled, if necessary, with available universal vascular cartridges (30 mm - 2.0 mm, EndoGIA™, Medtronic) as the supply of green cartridges was limited. The division exposes the bronchial bifurcation between the upper and the lower lobar bronchus. The lobar bronchi are alternatively clamped to allow individual ventilation of each lobe, reproducing the conditions of lobectomy.

### 2.5. DIC parameters

DIC was performed simultaneously with the ventilation tests (section 2.3). The VIC3D commercial software suite (version 7.2.4, Correlated Solutions Inc.) analyzed a series of image using a *subset-based* approach. The density and average dot size of the speckle pattern restricted the size of the subset and therefore the accuracy and resolution of the measurements. The best subset size (Table 1) was calculated by the software. To do so, it digitally applied a known strain with noise to the reference image of the sample. It then scanned the deformed image with different window size, called subset size, until the calculated strain on the deformed image and the known strain applied to the reference image matched. This subset size minimized the correlation error. This correlation error (mean of 0.0101 pixel over all samples) is used to determine if the measured displacement values, or computed strain values, are valid. The correlation error values on the edges of the lung do not appear to be higher than the ones further away from the edge of the specimen. Thus, the presence of strain peaks does not appear to be connected to the reliability of the DIC method (Supplementary materials).

**Table 1:** DIC parameters. Mean and standard deviation (SD) of the subset size, step and spatial resolution of displacement and of deformation for the analyzed samples. SIZE: 2 COLUMNS-FITTING

| DIC<br>Param. | Resolution (mm) |  | Step<br>(px) | Strain filter<br>(data points) | Sr<sup>d</sup> = Subset (px) |  |  | Sr<sup>e</sup> (px) |  |  | AOI (px<sup>2</sup>) |  |  |
| --- | --- | --- | --- | --- | --- | --- | --- | --- | --- | --- | --- | --- | --- |
|  | x | y |  |  | *healthy* | *inferior* | *superior* | *healthy* | *inferior* | *superior* | *healthy* | *inferior* | *superior* |
| **Mean** | 0,155 | 0,155 | 2 | 15 | 61,4 | 67,2 | 63,5 | 89,4 | 926,3 | 1004,1 | 234568 | 146542 | 86512 |
| **SD** | 0,004 | 0,003 | 0 | 0 | 9,6 | 14,5 | 12,5 | 9,6 | 14,5 | 12,5 | 51169 | 30246 | 21376 |

The software interpolates the displacement of the pixels in between the analyzed points. The *step size* is the space in pixels between two data points analyzed during correlation. To achieve a compromise between the computational time and the smoothing effects due to interpolation, the analysis used a step size of 2 (Table 1). An optimized 8-tap interpolation scheme measured the sub-pixels displacement. The Lagrangian strain computation used a *filter size* window of 15 data points to smooth the results, which represents an area of 30 pixels (step*filter size). The filter size was chosen inferior to 1/3^rd^ of the subset size to avoid over-smoothing the computed strains. The distance between two independent displacement measurements defines the displacement spatial resolution Sr^d^ while the distance between two independent measurements of strain measurements defines the strain spatial resolution Sr^ε^ calculated as Step × (Strain Filter-1) + Subset. The noise of the DIC measurement depends on the cameras focus and lenses, lighting, glare, angle between cameras, subset size and speckle pattern quality. The noise level is evaluated as the mean variation of displacement or strain of the AOI at a resting state (unloaded specimen) at ± 25 µm on the displacement measurement and of ± 0.06 % on the strain measurement. The noise values are significantly lower than the measured displacements and strains (strain peaks are 2.5 to 73.8 times higher than the noise). A custom algorithm [31] was developed in Python (version 3.0) to reproduce all results obtained by DIC [30].

The cameras capture the 3D area of interest (AOI, Table 1) representing the whole lung of approximately 234568 ± 51169 px^2^ (mean ± sd) with a pixel resolution of about (0.155 x 0.155) ± (0.004 x 0.003) mm^2^/px^2^. The average subset size on the performed experiments is 64 ± 12 pixels (∼10 mm, Table 1) and represents about 1/10^th^ of the length of the staple line (∼120 mm, section 2.4).

### 2.6. Compliance evaluation and statistical analysis

The lung compliance represents the slope of the linear portion of the pressure-volume curve [32]. As the portion of the pressure-volume inflation curves was not linear before 5 cmH_2_O, only pressure data above 5 cmH_2_O was used for the compliance calculation. Also, since the system took about two cycles to reach a stable volume, data after the 3^rd^ cycle was used in the compliance calculation. As the compliance is measured clinically during a volume intake, only inflation portions of the data were kept. The pressure-volume curves were plotted from raw volume data and interpolated pressure data obtained after kriging of the raw pressure data. A custom algorithm [30] was developed in MATLAB (version R2018b) to calculate the lung compliance: the slope of the linear regression of pressure-volume curves presented above, without considering data drifting due to leakage. If the maximum volume data of the considered cycle was 5% above the maximum volume of the 3^rd^ cycle, then data of that cycle was removed as considered drifting due to a leak. On Python 3.0, a right tailed paired Wilconxon sign test (with no continuity correction) allowed statistical comparison (α = 5 %) of lung compliance before and after resection on a sample population of 9 specimens. Specimen #1, #5 were eventually removed from this analysis as they leaked at a healthy state or presented extreme volume values. On Python 3.0, a left tailed paired Wilcoxon sign test (with no continuity correction) allowed statistical comparison (α = 5 %) of lobes burst pressure with sample #2 burst pressure approximated to 60 cmH2O to confirm the statistical inequality shown in clinical studies.

## 3. Results

This section focuses first on a detailed visual description of the air leak obtained after pulmonary resection and then on a quantitative description of the physiological parameters, volume, pressure and strain, influencing the leak.

### 3.1. Physiology of the leak

Observation of the lung tissue provides a visual characterization of the leak (Figure 3). First, the stapler seals the lungs tightly at the cutting line, where no air bubbles are observed (Figure 3–B). Then, a dark red coloring appears along the staple line (Figure 3– C). The lung tissue is bruised: it bleeds and the blood spreads under the pleura suggesting that stapling locally traumatizes lung tissue. Generally (7/8 specimens, Table 2), leaks occur at the perforation of the pleura and lung tissue made by the staple legs (Figure 3A) as shown in Figure 3-D, E. The eighth specimen (specimen #2, inferior lobe) showed a leak from a tear in the pleura, not related to stapling but probably due to mishandling of the tissue (Table 2). The pleura does not retract from the cut-line to expose the lung tissue: the intact pleura is easily recognizable between the staple rows as a smooth glistening membrane covering the lung surface (Figure 3–F). The pleura of the previously cited specimens tears locally, from a staple hole of the inner row (furthest from the cutting line), over a length of several staples, exposing the lung tissue (Figure 3–E). The observation of the leak corresponds to the corner between the torn pleura and the staple hole (Figure 3–D, E), the tissue portion on the costal surface of the staple line leaked (8/8 specimens, Table 2).

**Table 2:** Experimental results of the burst pressure (cmH_2_O), visual location of the leak and the quality of the staple line for all the tested specimens. SIZE: 1.5 COLUMN-FITTING

| Specimen | P burst (cmH<sub>2</sub>O) |  | Location of the leak |  | Quality of SL |
| --- | --- | --- | --- | --- | --- |
|  | *inferior* | *superior* | *inferior* | *superior* |  |
| 1 | 5 | 5 | c, m | c, ne | *** |
| 2 | 10 | >60 | c, ne | no leak | *** |
| 3 | 30 | 30 | c, e | c, m | ** |
| 4 | 7 | 25 | c, m | c, ne | ** |
| 5 | 5 | 50 | c, m | c, ne | *** |
| 6 | 10 | 10 | c, m | c, ne | *** |
| 7 | 25 | 20 | c, m, ne | c, ne | *** |
| 8 | 45 | 29 | c, m | c, m | ** |
| 9 | 10 | 45 | N.A. | N.A. | *** |
| 10 | 10 | 60 | N.A. | N.A. | ** |
| 11 | 10 | 7 | N.A. | N.A. | *** |
| **Mean** | 15.2 | 31.0 |  |  |  |

N.A. Not Available
SL: Staple Line
c = costal side
m = middle of SL
ne = near-end of SL
\* poor
\*\* acceptable
\*\*\* good

**Figure 3:**
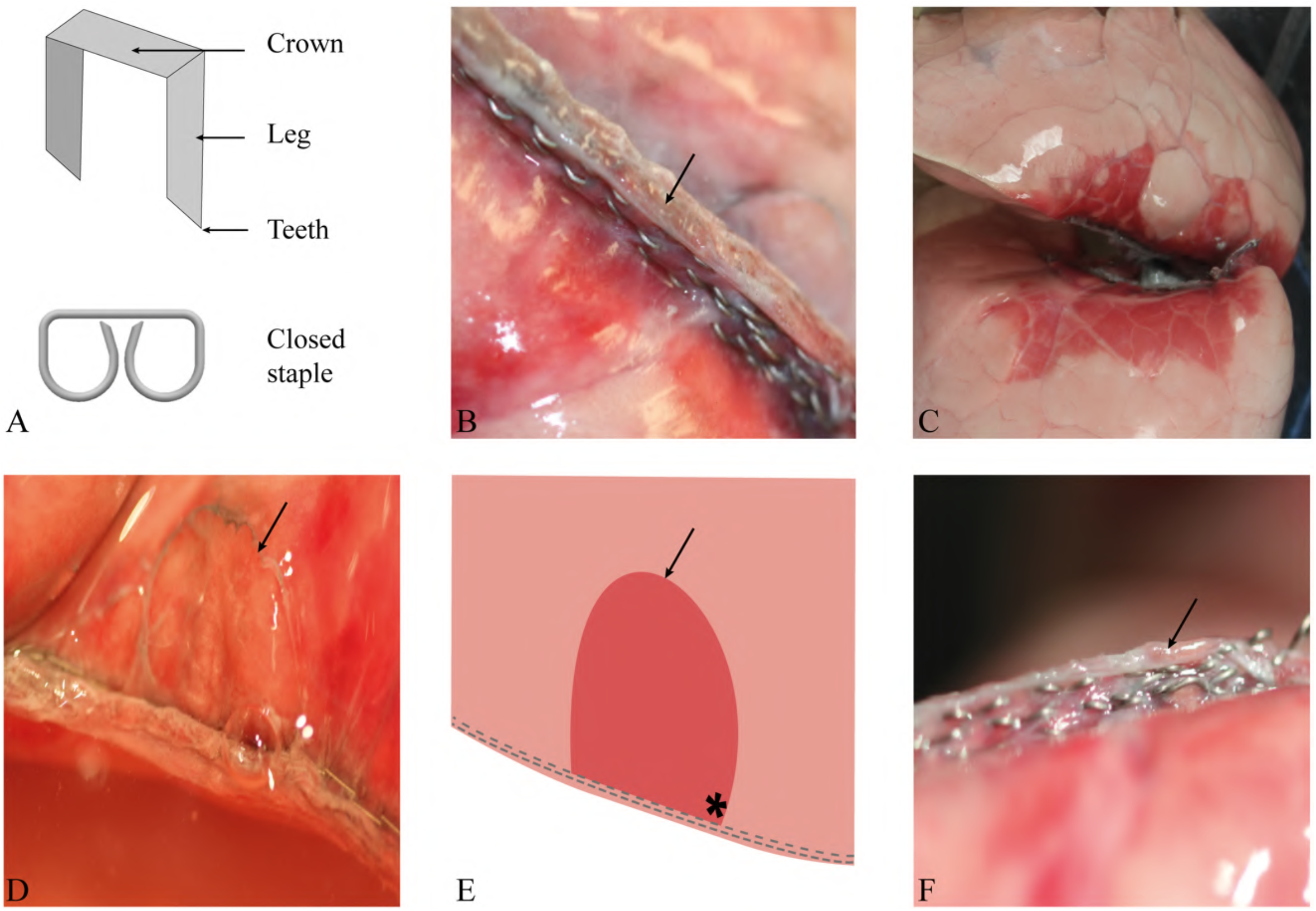
Visualization of an air leak on a stapled lung. A – Nomenclature of a staple shown before and after stapling. B – Exposed lung tissue at the cut-line, sandwiched in between the two visceral pleura by the stapling mechanism. Arrow points to the lung tissue at the cut line. C – Trauma of the lung tissue localized around the staple line. D – Air bubble leaking at staple hole near the exposed lung parenchyma because of a torn visceral pleura. Arrow shows tearing contour of the visceral pleura. E – 2D schematic of picture D. Arrow points to the torn visceral pleura exposing the lung tissue. Star sign indicates the leak site. F – Bulge of pleura near the cut-line indicated by arrow. SIZE: 1.5 COLUMNS-FITTING

### 3.2. Compliance

After examining where the leak develops, this subsection focuses on whether the lung tissue reacts differently in terms of pressure and volume depending on its state: healthy (entire lung) or stapled (superior or inferior).

If the mean Area Of Interest (AOI, Table 1) of the entire lung represents 100% (234568 px^2^), the lower lobe therefore accounts for 2/3rds (146542 px^2^) of the total lung area compared to 1/3rd for the upper lobe (86512 px^2^). The bursting pressure (Table 2, Figure 4f) represents the pressure at which the inferior or superior lobe leaked during the experiment. The lower lobe statistically leaked at a smaller burst pressure (*p-* value<0.046, Figure 4f) than the upper lobe.

**Figure 4:**
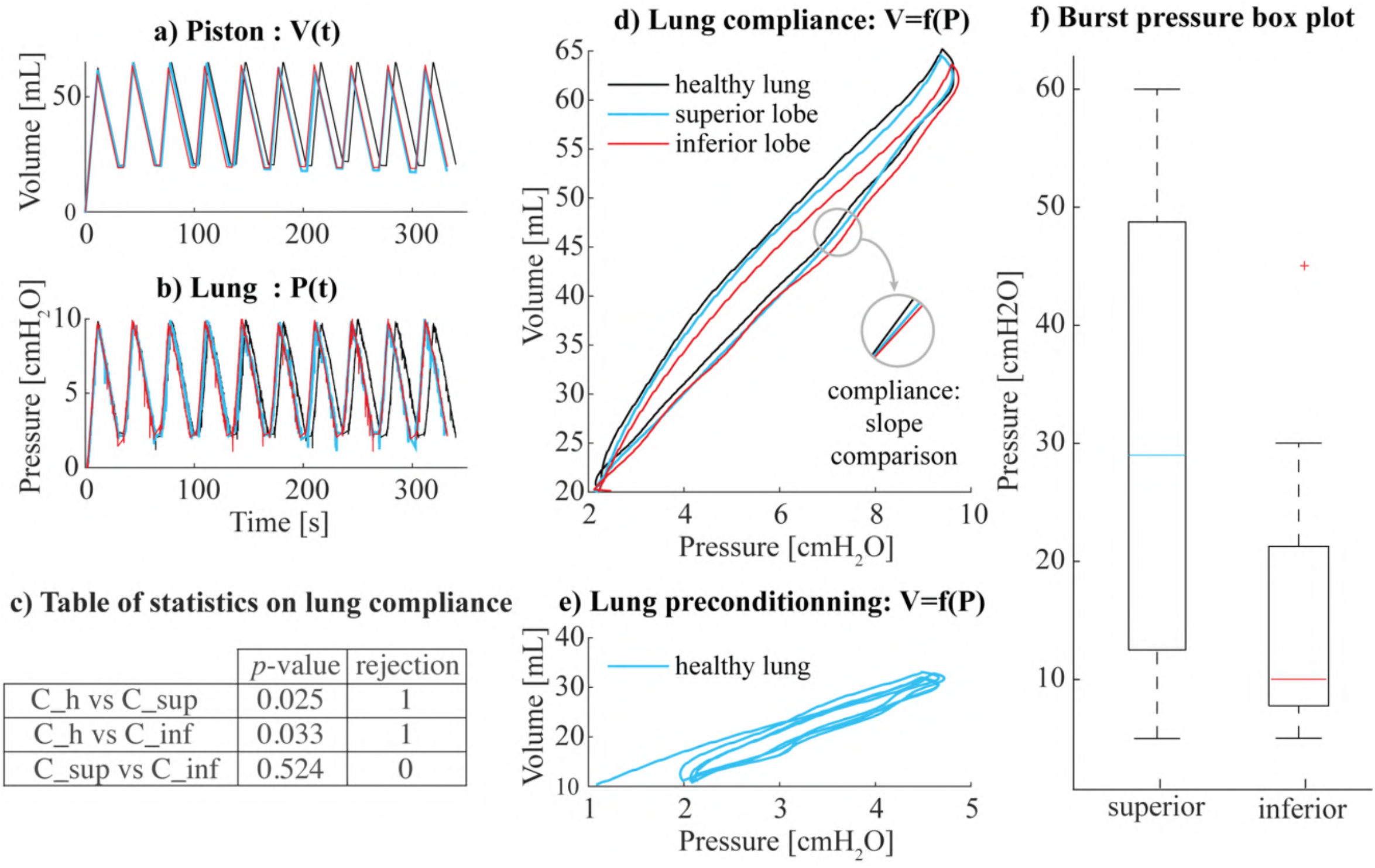
Typical experimental results (specimen #7). a) Volume (mL) delivered by the piston to the lung over time (s); b) corresponding lung pressure (cmH_2_O) over time (s); c) table of statistics on lung compliance results; d) lung compliance hysteresis curves: pressure (after kriging) against volume curves of the 3rd cycle (to avoid plot overload), for the healthy lung (black), superior (blue) and inferior lobes (red); e) plot of the lung compliance curves of the odd cycles for the healthy lung preconditioning; f) burst pressure box plot of inferior and superior lobes. SIZE: 2 COLUMNS-FITTING

One may wonder why the larger lower lobe leaks more easily than the smaller upper lobe if both lobes are considered to have the same mechanical properties and elasticity. Compliance assessing the relationship between pressure and volume is therefore an interesting indicator to complement this result. The lung compliance evaluates the ability of the lung to deform in response to a variation of pressure. Comparing healthy vs inferior/superior compliances quantifies the effect of pulmonary resection on the lung tissue (Figure 4, Table 3). Figure 4d shows three representative pressure-volume curves, one for each state, based on smoothed discrete measurements as explained section 2.6. First, compliance curves present a hysteresis behavior: the inflation curve is lower than the expiration curve. Secondly, the preconditioning led to the overlap of compliance curves after the 5^th^ cycle, confirming that there are no leaks in the system. Mean compliance differences are respectively of 9.3% and 10% between normal lung vs superior/inferior lobes (Table 3). Finally, the compliance of the healthy lung is statistically superior to that of the upper lobe (*p*<0,025, Figure 4) and of the inferior lobe (*p*<0,033, Figure 4). However, there was no significant difference between lobar compliances (*p*<0,524, Figure 4).

**Table 3:**
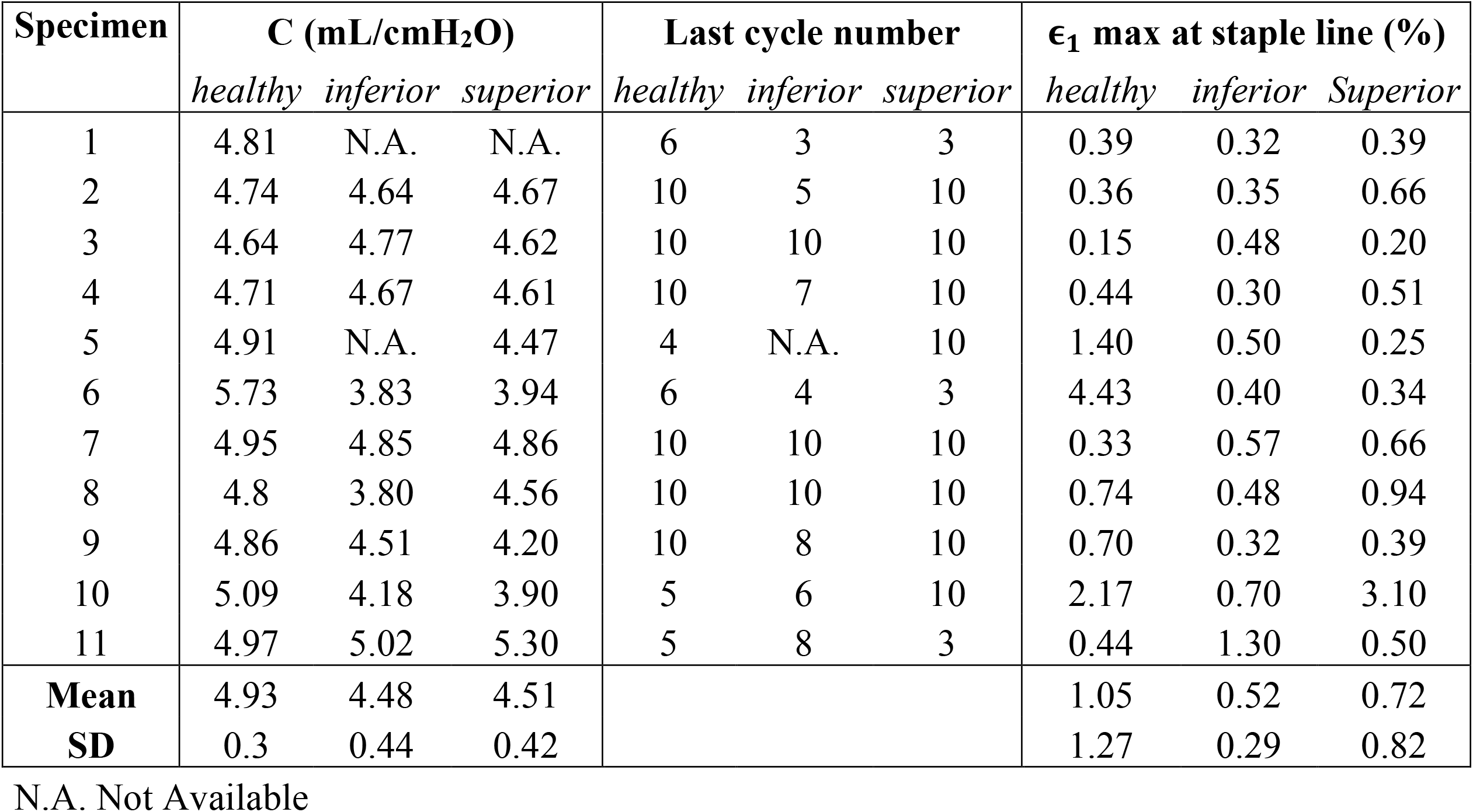
Experimental ventilation and strain data for the healthy lung, as well as for inferior and superior lobes: compliance C (mL/cmH_2_O), last cycle number used in the compliance calculation, maximum principal strain ϵ_1_ (%) at staple line calculated with DIC at the maximum pressure of the last cycle before leakage. SIZE: 2 COLUMS-FITTING

| Specimen | C (mL/cmH<sub>2</sub>O) | | | Last cycle number | | | $\epsilon_1$ max at staple line (%) | | |
| --- | --- | --- | --- | --- | --- | --- | --- | --- | --- |
|  | *healthy* | *inferior* | *superior* | *healthy* | *inferior* | *superior* | *healthy* | *inferior* | *Superior* |
| 1 | 4.81 | N.A. | N.A. | 6 | 3 | 3 | 0.39 | 0.32 | 0.39 |
| 2 | 4.74 | 4.64 | 4.67 | 10 | 5 | 10 | 0.36 | 0.35 | 0.66 |
| 3 | 4.64 | 4.77 | 4.62 | 10 | 10 | 10 | 0.15 | 0.48 | 0.20 |
| 4 | 4.71 | 4.67 | 4.61 | 10 | 7 | 10 | 0.44 | 0.30 | 0.51 |
| 5 | 4.91 | N.A. | 4.47 | 4 | N.A. | 10 | 1.40 | 0.50 | 0.25 |
| 6 | 5.73 | 3.83 | 3.94 | 6 | 4 | 3 | 4.43 | 0.40 | 0.34 |
| 7 | 4.95 | 4.85 | 4.86 | 10 | 10 | 10 | 0.33 | 0.57 | 0.66 |
| 8 | 4.8 | 3.80 | 4.56 | 10 | 10 | 10 | 0.74 | 0.48 | 0.94 |
| 9 | 4.86 | 4.51 | 4.20 | 10 | 8 | 10 | 0.70 | 0.32 | 0.39 |
| 10 | 5.09 | 4.18 | 3.90 | 5 | 6 | 10 | 2.17 | 0.70 | 3.10 |
| 11 | 4.97 | 5.02 | 5.30 | 5 | 8 | 3 | 0.44 | 1.30 | 0.50 |
| **Mean** | 4.93 | 4.48 | 4.51 |  |  |  | 1.05 | 0.52 | 0.72 |
| **SD** | 0.3 | 0.44 | 0.42 |  |  |  | 1.27 | 0.29 | 0.82 |

N.A. Not Available

### 3.3. Strain field analysis

This subsection first describes the biomechanics of the healthy lung, inferior lobe and superior lobe. Then, the location of maximum major principal and shear strains on the resected specimens obtained by DIC are compared to the observed leaking area to determine a possible co-localization between the two.

The DIC measures the strain of the lung during the experiment at different pressures (Figure 5). Especially, major principal strain (*ϵ*_1_) corresponding to the maximum strain and in-plane shear strain *γ*_*max*_ = (*ϵ*_1_ − *ϵ*_2_) for each analyzed point. The major principal strain is increasing with pressure as the lung inflates. The lung edges inflate first as the principal strain *ϵ*_1_ is higher on the edge areas (>0.6%) versus in central areas near the air supply tube (∼0.1%). The shear strain is also higher on the edges. For each state of the lung, the shear strain is not equal to zero (>0.1%) meaning the principal strains are not equal: lung does not inflate like a party balloon which stretches under equibiaxial strain (Figure 5). Note that on Figure 5, a smoothing filter hided the small scale variations in the strain fields to highlight the whole lung scale mechanics.

**Figure 5:**
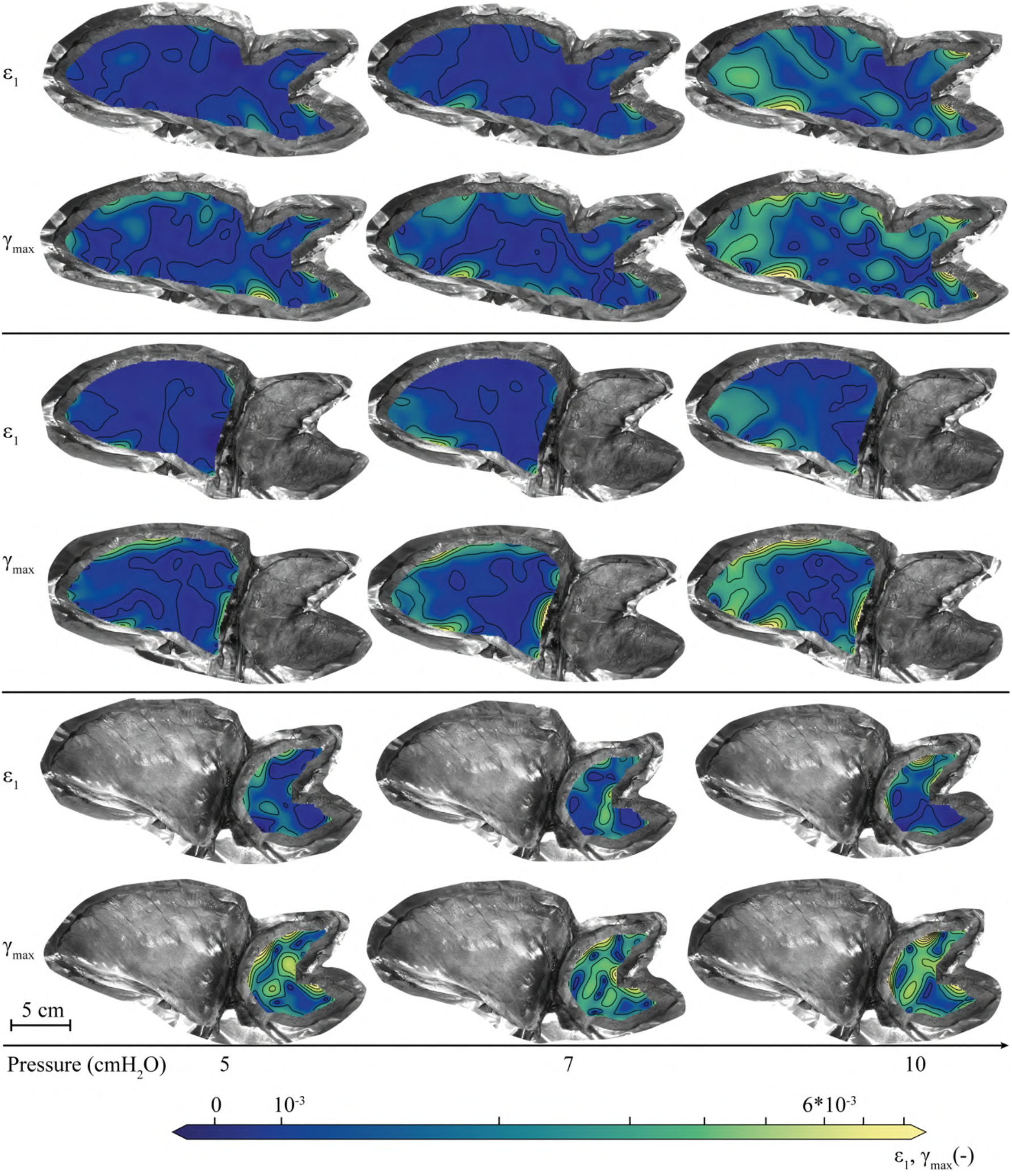
Time lapse of smoothed major principal (ϵ_1_) and shear (*γ*_*max*_) strains developing in the healthy lung (top panel) as well as in its inferior lobe (central panel), and its *superior* lobe (bottom panel) for different pressure levels (specimen #3). The strain patterns were smoothed with a subset of 101 px, a strain filter of 15 and a step of 7. SIZE: 2 COLUMNS-FITTING

After DIC measurements, the immersed inflated lung presented air bubbles at specific locations noted on a 2D schematic in Figure 6 A. The leak site was along the staple line for 7/8 samples (1/8 did not present leaks but pleura detachment). Among these, the air leak happened at the center of the staple line for the lower lobe and at the near the posterior end of the staple line for the upper lobe in 5/8 cases. The leaks were also only observed along the costal surface of the staple line for all leaking specimens (Figure 6 A, Table 3).

**Figure 6:**
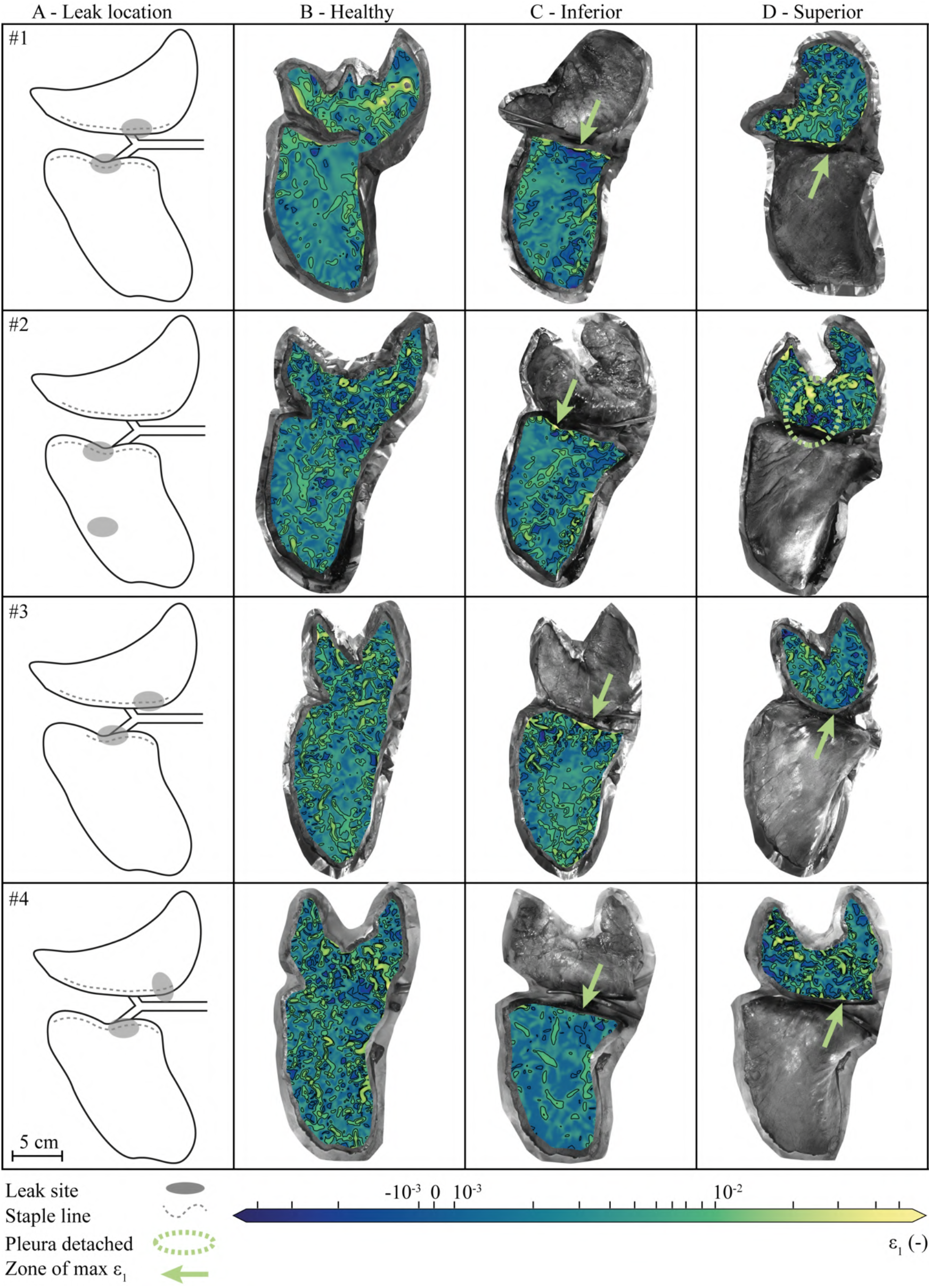
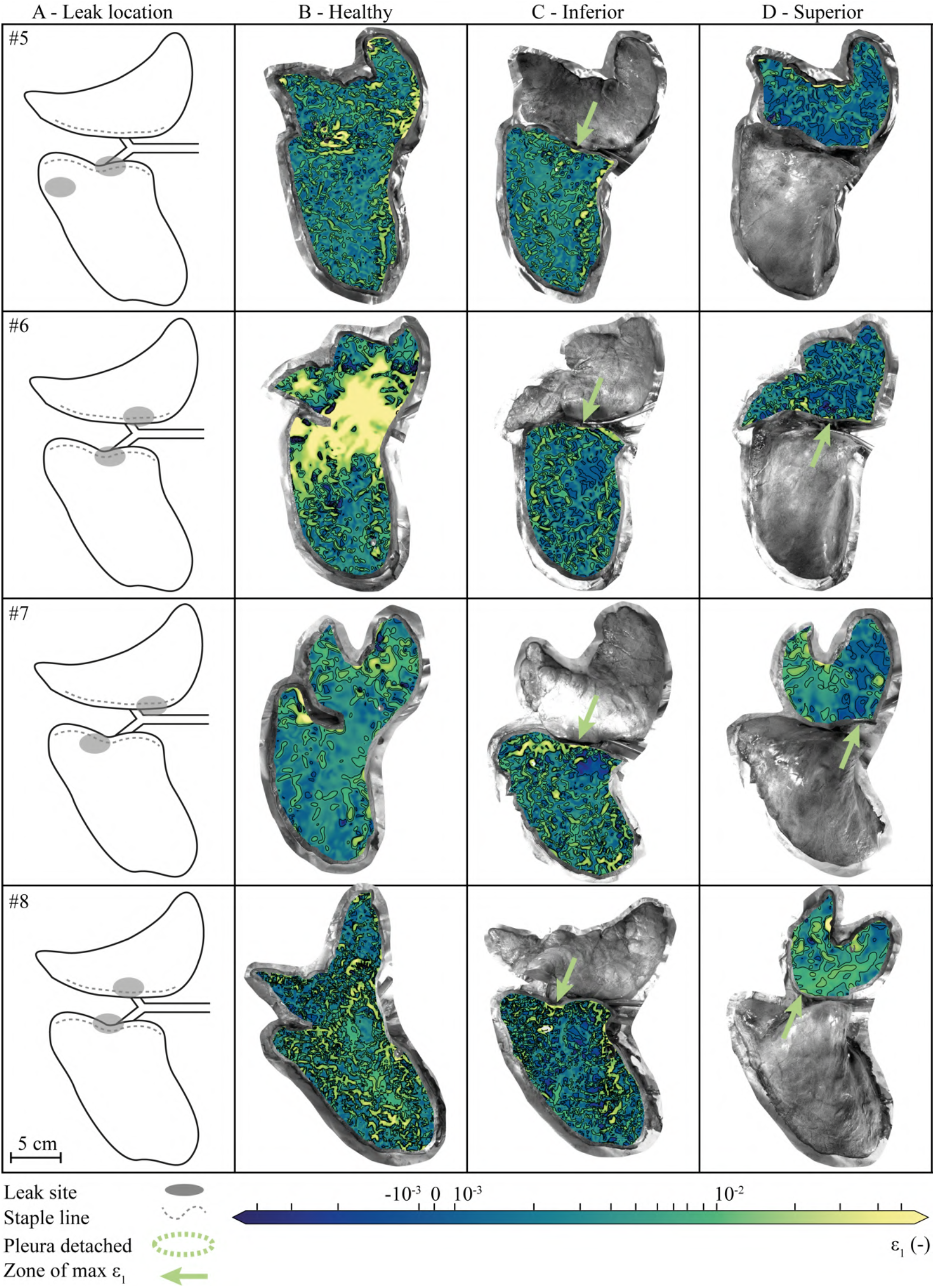
Leak sites and lung principal strain (*ϵ*_1_) patterns. Schematic localization of observed leak site on the porcine left lung (A). Principal strain contour plots for the entire lung (B), inferior (C) or superior lobe (D) independently inflated with green arrows pointing to zones of maximum major principal strain at staple line. The same symmetrical logarithmic principal strain scale is used for all specimens. SIZE: 2 COLUMNS-FITTING

A comparative analysis between the strain patterns of the healthy lung and those of the resected lobes was carried out to study the biomechanical interaction of staples with the lung tissue. For each specimen, the principal strain (*ϵ*_1_) patterns are plotted for its three states (healthy lung, superior lobe, inferior lobe) at 10 cmH_2_O and presented using the same strain scale (Figure 6 B, C and D). The major principal strain (*ϵ*_1_) scale ranges from less than -0.1%, representing tissue contraction, to more than 1%, representing tissue elongation. The maximum principal strain is on average 1.05%, 0.52%, 0.72% at 10 cmH_2_O of pressure for the healthy lung, inferior lobe and superior lobe respectively (Table 3). The same comparative analysis is conducted for the shear strain (*γ*_*max*_) patterns (Figure 7). Specimen #5 (Figure 7D) did not exhibit a leak and showed a uniform distribution of shear strain along the staple line contrary to leaking specimen, which have shear strain concentration superior to 1% at the staple line (ex: #2 Figure 7C). The zone of maximum major principal strain and shear strain at staple line are indicated with green arrows and corresponds to the observed leak location (Figure 6, Figure 7).

**Figure 7:**
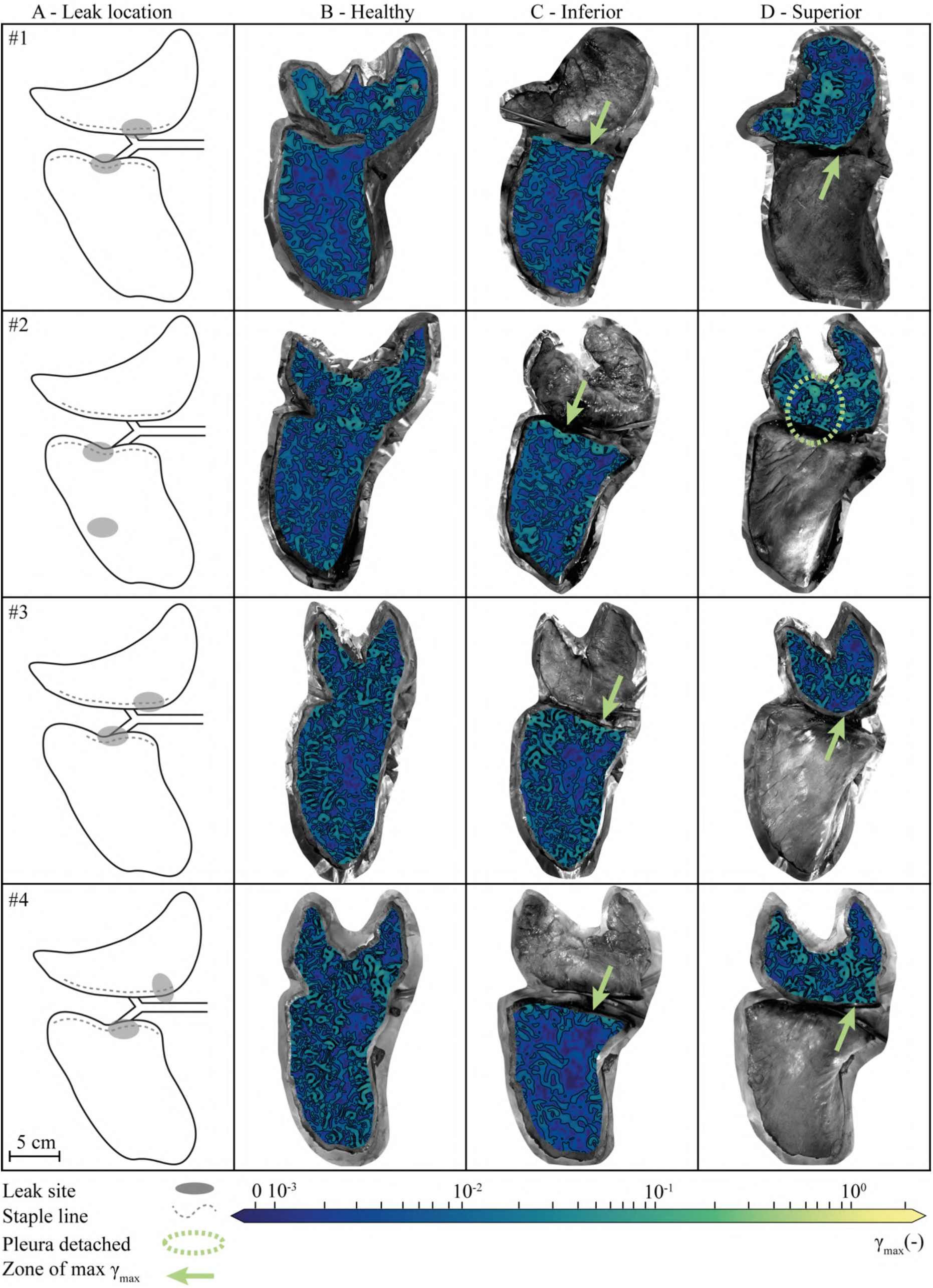
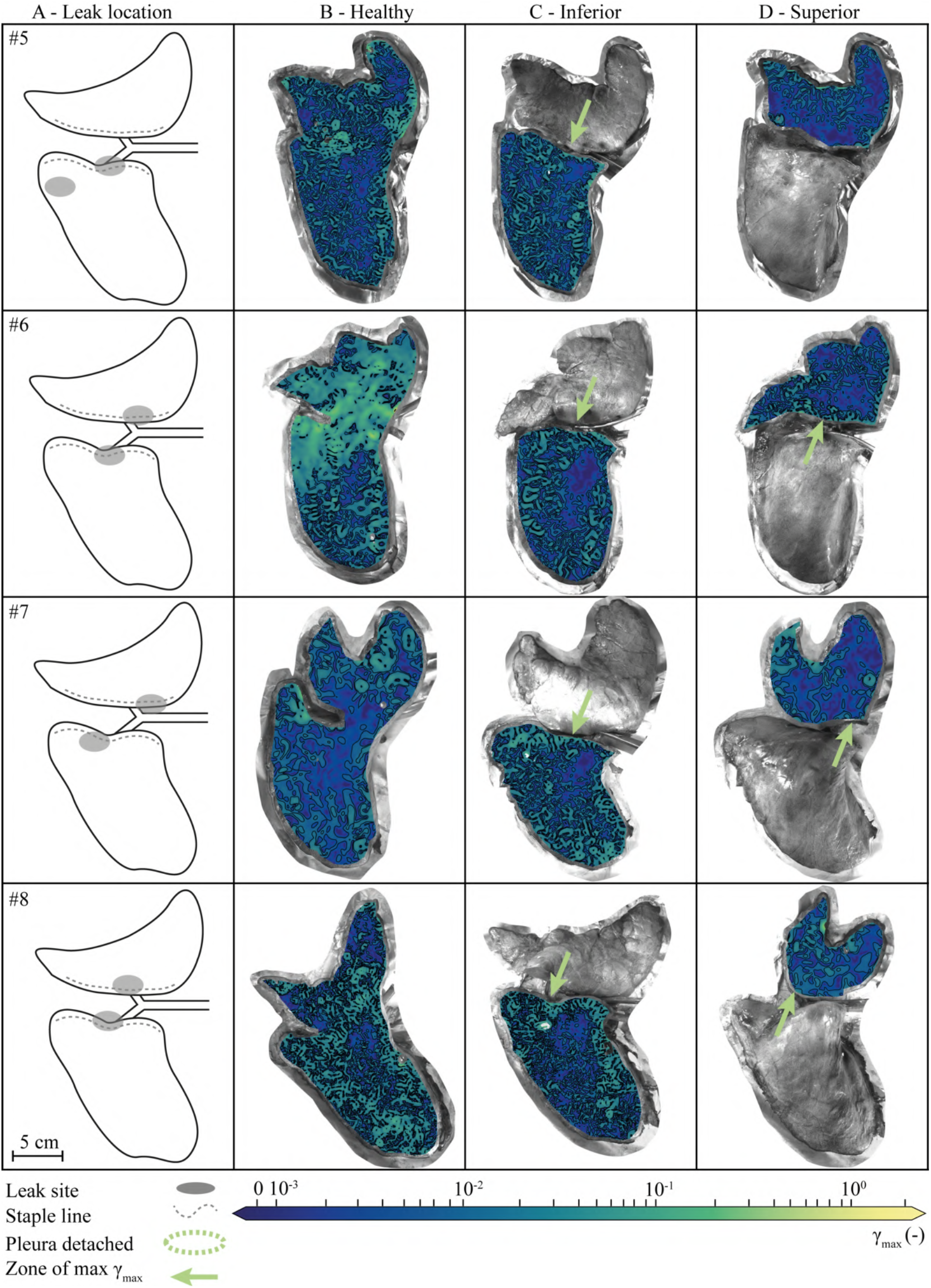
Leak sites and lung maximum shear strain (*γ* = *ϵ*_1_ − *ϵ*_2_) patterns. Schematic localization of observed leak site on the porcine left lung (A). Shear strain contour plots for the entire lung (B), inferior (C) or superior lobe (D) independently inflated with green arrows pointing to zones of maximum shear strain at staple line. The same symmetrical logarithmic maximum shear scale is used for all specimens. SIZE: 2 COLUMNS-FITTING

## 4. Discussion

In this study, lung tissues with and without staples were biomechanically and visually characterized in order to gain knowledge on air leakage following pulmonary resection. This discussion is meant to provide new insights on potential relationships between leak sites, lung compliance and strains development in healthy lung as well as in its superior and inferior lobes following stapling. These results could be used as a basis for future investigations on how to minimize air leakage by changing the stapling devices or techniques.

### 4.1. Air leaks at staple holes of inner row from exposed lung tissue due to torn pleura, on the costal side

Although this result seems intuitive, this visual characterization of the leak (Figure 3) is evidence to support that the cut line is not leaking. This first observation shows that the stapling technique provides an airtight tissue boundary despite the variation in stapled tissue thickness and uniform cartridge size choice. The visceral pleura is locally torn around the staple holes from the inner row of the staple line and these holes present leak because the lung tissue is exposed and traumatized. The costal surface is identified as the most vulnerable to leakage. The observation of a leak on the costal surface is consistent with anatomy since this surface inflates most even in physiological conditions. Indeed, the mediastinal part of the lung is anatomically constrained by mediastinal structures such as the heart [33], inducing less deformation to the mediastinal lung tissue. In our experimental conditions, the weight of the lung constrained the mediastinal surface against the table replicating the physiological conditions. However, no test was performed by turning the stapler or the lung over to ensure that these parameters do not impact this result.

Regarding the leak observation sites, the presented results are generally consistent with the literature: leaks occur at staple lines. However, the study of Imhoff *et. al*. reports leakage at extremities of the staple line [7], [8]. In our experiment, leakage is observed in the center of the staple line for the lower lobe and near the posterior extremity end for the upper lobe (Table 2). If the leak was a result of any technical defect, one would have expected that each lobe would have been affected in a similar way on both side of the staple line, although this should be confirmed on a larger sample size. The leak location being at the center of the staple line could be due to anatomical reason as it is the location of the bronchus insertion. The leak location could also correspond the location of the application of the second staple cartridge and possible overlaps or misalignments (Figure 8a) or misconnections between successive staple lines (Figure 8b) of the staple line. At the extremities it may be that the last staples do not always cling well (Figure 8b), not always find lung tissue for both legs to cling to, reducing their effectiveness. To avoid these technical issues, a thoracic surgeon trained the experimenter to obtain accurate and clean staple lines. Once the lung tissue was stapled, the quality of the staple line was visually assessed by the experimenter using a scoring system (Figure 8). No specimen was rejected for its score. This score was not able to identify a relationship between the leakage and a technical problem. This study provided concrete evidence on the location and origin of air leaks as a knowledge base for further research on post-operative air leaks.

**Figure 8:**
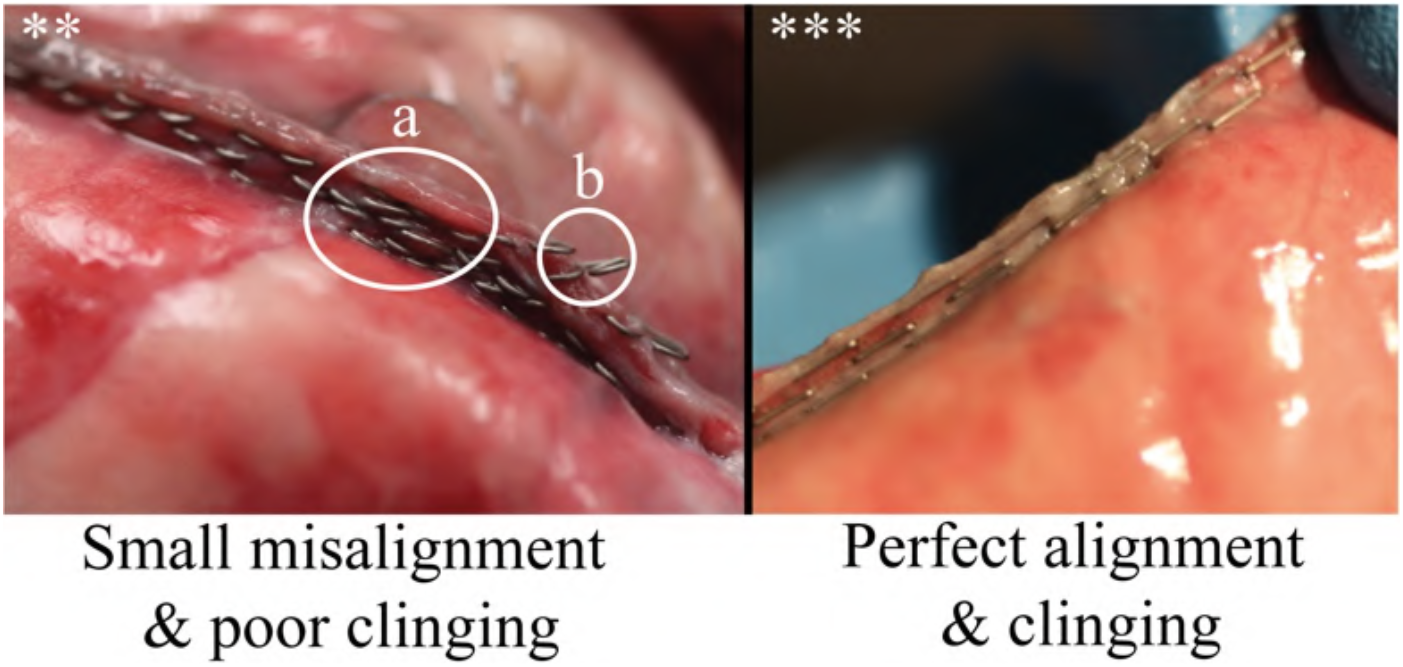
Staple-line nomenclature and scoring system: *Poor: very poor clinging or alignment combined with misconnections of the staple line (no specimen), **Acceptable: (a) small misalignment or (b) poor clinging of the staple line (4/11), ***Good: perfect alignment and clinging of the staple line (7/11). SIZE: 1 COLUMNS-FITTING

### 4.2. Observed leak site corresponds to zones of maximum principal strain at the staple line

To investigate the localization of leak at the staple line, one can visually compare the staple line leak sites on each specimen (Figure 6A) with the sites where the maximum principal strain is concentrated at the staple line (Figure 6C, D, green arrows). Best example: specimen #5D exhibits a uniform distribution of the principal strain along the staple line and one concentration point of principal strain that turned out to be the same location as its leak. Moreover, two specimen (specimen 5 & 6, Figure 6) globally exhibited higher principal strains (ϵ_1_) at healthy state, which is compatible with the relatively high compliance (more flexible to deformation) of the two specimens (4.912 mL/cmH_2_O & 5.726 mL/cmH_2_O compared to an average of 4.774 mL/cmH_2_O for the other specimens, Table 3). For each lobe, this trend repeats: the major strain concentration occurred at the leak observation site (Figure 6C, D). However, more than one area of maximum principal strain can be identified on the strain pattern of each specimen but not all of them were reported to be leak sites (Figure 6 #4D). On specimen #7D, the arrow supposedly points to the maximum strain area corresponding to the leak site but there is no visible maximum strain. Furthermore, principal strain concentrations (Figure 6) seem to be a better indicator of air leak location than maximum shear (Figure 7). The DIC technique requires several measurement points (almost half of a subset) to compute the strain. This is why the strain contour plot is smaller than the photographed lung, which could explain the unseen point of maximum principal strain concentration on specimen #7D. Maximum major principal strain area can also be explained by a detached visceral pleura near the staple line as seen in specimen #2D. Another limitation could be the position of the staple line relatively close to the air input however the lobar bronchus delivers air first to the lung edges as shown by the higher major principal strain at the edges (Figure 5). As the lung was checked for misplaced tubing in the main bronchus, it should have no apparent reason to leak except for a torn visceral pleura near the staple holes. This study compared results of major strain site with leak observation site and cannot conclude to a causal relationship, perhaps further research could focus on investigating this causal relationship.

### 4.3. Left upper or lower lobe resections have similar compliance results but lower than healthy lung

There is a minimal change in compliance between whole lung and individual lobes (Table 3). Indeed, healthy lung compliance decreases to 91% of its prior value after upper or lower resection. There was no statistical difference between the compliance of the left upper lobe and left lower lobe. Studies reported a proportional decrease in respiratory system compliance relative to the amount of resected tissue assuming no change in the chest wall compliance, which means that the lung compliance would follow the trend of the respiratory system compliance [19]–[21]. The difference between lobes compliance and healthy lung compliance is minimal but significant. One might have expected that lobe compliances would have decreased with respect to the volume proportion of the lobes compared to the entire lung. This result is counterintuitive and not completely consistent with the conclusions of previous studies [19]–[21]. Our experiments were not designed to assess why this would be the case, but it is reasonable to infer that it may be related to internal pulmonary architecture or alveolar ultrastructure. The alveoli are more numerous and smaller in the lower lobe and less numerous and larger in the upper lobe, which can complicate interpretations of compliance results. The mechanical behavior of the healthy lung was assessed only on the basis of low pressure and therefore low deformation values. This behavior could change for larger deformation values and involve other response mechanisms to pressure. The elastin fibers that compose the lung tissue support the efforts during low deformations while the collagen fibers support the efforts for higher deformations [34]. This study provides new data to the literature on changes in lung compliance at low pressures in positive-pressure ventilation. Further studies should aim to validate these results and investigate how compliance is related to strain patterns at different ventilation pressures. This could influence the pressure practices applied to post-operative chest drains to avoid imposing too much pressure that would imply a strong change in compliance.

### 4.4. Compared to upper lobes, lower lobes statistically burst at lower pressures in *ex vivo* conditions

This experiment measured the burst pressure of both left upper and lower lobes (Table 2, Figure 4f). The left lower lobe statistically leaked at lower pressures (*p-*value<0.046). In several studies, upper lobectomy, which correspond to the remaining of the lower lobe, was associated with a higher risk of developing a leak [16], [17], [35], [36]. More specifically, Casha *et. al*. conducted a finite element analysis on a simplified lung model and suggested that the lower lobe is more prone to leakage because it has to reshape itself to fit the bullet shape of the ribcage [1]. However, with the present results, the conclusion that the lower lobe of the left lung leaks at lower pressures is not dependent on the rib cage since the study was conducted *ex vivo*. Rather, this difference in leak burst pressures may be related to the unequal distribution of overall strains along staple lines in the superior and inferior lobes. One specimen did not leak and exhibited a more uniform distribution of maximum shear strain and major principal strain. If the visceral perforation is considered as a crack there may be a causal relationship between the development of air leaks and the major principal strain patterns observed. Taken together, these results suggest that strain patterns are crucial to understanding the mechanisms of staple line disruption and should be the focus of further investigation.

### 4.5. Limitations

The experiment has some methodological limitations. Porcine lungs may have a different mechanical behavior from that of humans. Pigs also have an additional lobe that does not exist in humans, which did not interfere in the dissection of the left lung. However, it is an animal model widely used in experimental studies seeking to understand lung biomechanics [37]. In addition, there were 8 valid specimens out of the 11 tested specimens, mainly due to missing information regarding the leak site, leading to a relatively small sample size. This study nevertheless provided statistical trends that could be compared with the literature. Finally, the experiment applied positive pressure insufflation, which does not represent physiological breathing but rather a physiological response to mechanical ventilation. The ventilation method took several precautions aiming at supplying air to the alveoli, however the resulting condition of the alveoli was not evaluated by imaging. In addition, lungs function within the physiological constraints of the chest cavity, trapped between the ribs and the diaphragm. In this experiment, the lungs were laid flat and horizontally in the open air rather than vertically and subjected to gravity under their own weight. These choices may have influenced the results. However, they were chosen to replicate the ventilation conditions of the lungs during surgery while giving visual access to the lung for DIC measurements. This is the first time that DIC was applied to the whole lung; this novel technique yielded valuable results for the lung biomechanics of stapling.

## 5. Conclusion

This experimental work describes with precision the physiology of air leakage following porcine lung resection. Air leaks originate from staple holes with a locally torn visceral pleura. They occur at the near end of the staple line of the upper lobe and at the center of the staple line for the lower lobe, both on the costal surface. The leak site is coupled with a concentration of major principal strain rather than a high shear strain developing at the staple line. This study opens perspectives that could lead to a practical focus on the stapled tissue region in order to minimize the stress concentrations associated with air leakage and the resulting clinical complications.

## Supplementary materials

Additional data associated with this article can be found in the online version at DOI: 10.5281/zenodo.4362643, type: dataset [30].

## Acknowledgment

The authors acknowledge helpful contributions and technical skills of team members from the laboratory for multiscale mechanics (LM2) as well as Maisonneuve-Rosemont Hospital’s laboratory manager Josée Tessier. This research was conducted as part of the activities of the TransMedTech Institute, thanks in part to the financial support of the Fonds de recherche du Québec and financial support of Mitacs (IT14660).

## Notes

http://doi.org/10.5281/zenodo.4362643

### Competing Interest Statement

This research has benefited from a surgical material donation from Medtronic, Canada. And George Rakovich receives speaker fees for Medtronic.

